# Homozygous c.259G>A variant in *ISCA1* is associated with a new multiple mitochondrial dysfunctions syndrome

**DOI:** 10.1101/089292

**Authors:** Anju Shukla, Malavika Hebbar, Anshika Srivastava, Rajagopal Kadavigere, Priyanka Upadhyai, Anil Kanthi, Oliver Brandau, Stephanie Bielas, Katta M Girisha

## Abstract

The iron-sulfur (Fe-S) cluster (ISC) biogenesis pathway is indispensable for many fundamental biological processes and pathogenic variations in genes encoding several components of the Fe-S biogenesis machinery, such as *NFU1, BOLA3, IBA57* and *ISCA2* are already implicated in causing four types of multiple mitochondrial dysfunctions syndromes (MMDS) among other human diseases. MMDS are clinically characterized by neurodevelopmental delay, neurological deterioration, lactic acidosis, extensive white matter abnormalities and early death. We report on two unrelated families, with two affected children each with neurodevelopmental delay, regression of developmental milestones, seizures, extensive white matter abnormalities, cortical migrational abnormalities, lactic acidosis and early demise. Exome sequencing of two affected individuals, one from each family, revealed a homozygous c.259G>A variant in *ISCA1* and Mendelian segregation was confirmed in both families. ISCA1 is a specialized factor known to mediate maturation of distinct Fe-S cluster (ISC) proteins. *In silico* functional analyses and structural modeling of the protein predict the identified *ISCA1* variant to be detrimental to protein stability and function. Notably the phenotype observed in all affected subjects with the *ISCA1* pathogenic variant is similar to that previously described in all 4 types of MMDS. The *ISCA1* variant lies in the only shared region of homozygosity between the two families suggesting the possibility of a founder effect. To the best of our knowledge this is the first instance where ISCA1 deficiency has been shown to be associated with a human disease, a new type of multiple mitochondrial dysfunctions syndrome.

---

Iron-sulfur (Fe-S) clusters (ISCs) are small inorganic cofactors of metalloproteins found in bacteria and eukaryotes that are indispensable for cellular processes such as respiration, protein translation, purine metabolism, DNA repair and gene expression regulation.^1-3^ The Fe-S protein biogenesis in eukaryotes occurs in mitochondria, cytosol, or nucleus and is carried out in two steps. The first step is the formation of [2Fe-2S] cluster by proteins forming the core ISC assembly components. This cluster is transferred to monothiol glutaredoxin 5 (GRX5), which acts as a Fe/S cluster transfer protein inserting the [2Fe-2S] cluster into mitochondrial [2Fe- 2S]-requiring proteins. The second step involves the generation of [4Fe-4S] components and their integration into the required metalloproteins (Figure S1A). Unlike the core ISC assembly components, proteins mediating the second step are not involved in formation of mitochondrial [2Fe-2S] and cytosolic [4Fe-4S] clusters.^4;5^ This delivery step requires highly conserved and functionally non-redundant A-type ISC proteins, ISCA1 and ISCA2 in eukaryotes.^6^ Loss of function of several Fe-S cluster assembly components have been known to result in several human diseases including multiple mitochondrial dysfunctions syndromes (MMDS), Freidreich’s ataxia, Swedish myopathy with exercise intolerance, and Sideroblastic anemia.^7-10^

Currently four types MMDS1 (MIM #605711), MMDS2 (MIM #614299), MMDS3 (MIM #615330) and MMDS4 (MIM #616370) have been mapped to *NFU1* (MIM #608100), *BOLA3* (MIM #613183), *IBA57* (MIM #615316) and *ISCA2* (MIM #615317) respectively. All MMDS share variable neurodevelopmental delay, regression, seizures, lactic acidosis and leukodystrophy resulting in early death of affected individuals.^11;12^ Here we report another likely MMDS resulting from a biallelic mutation in *ISCA1*, a key component of the Fe-S biogenesis process, from two unrelated Indian families.

Family 1 consulted us for prenatal counseling. They had two healthy daughters and lost two children (Figure 1A). Both II.3 and II.4 had normal birth and antenatal history followed by inconsolable cry and feeding difficulties since newborn period. They had progressive neurological deterioration after a brief period of normal growth. II.3 died at 1 year and 7 months while II.4 succumbed at 5 years of age. Stored DNA was obtained for patient II.4 and blood samples were obtained for parents and healthy children. Brain imaging of both children showed pachygyria, extensive cerebral and cerebellar white matter disease and dilated cerebral ventricles. Increased lipid lactate peak was seen on magnetic resonance spectroscopy of brain in II.3. Normal hematological and biochemical investigations were noted except for elevated creatinine phosphokinase in II.4.

**Figure 1:**
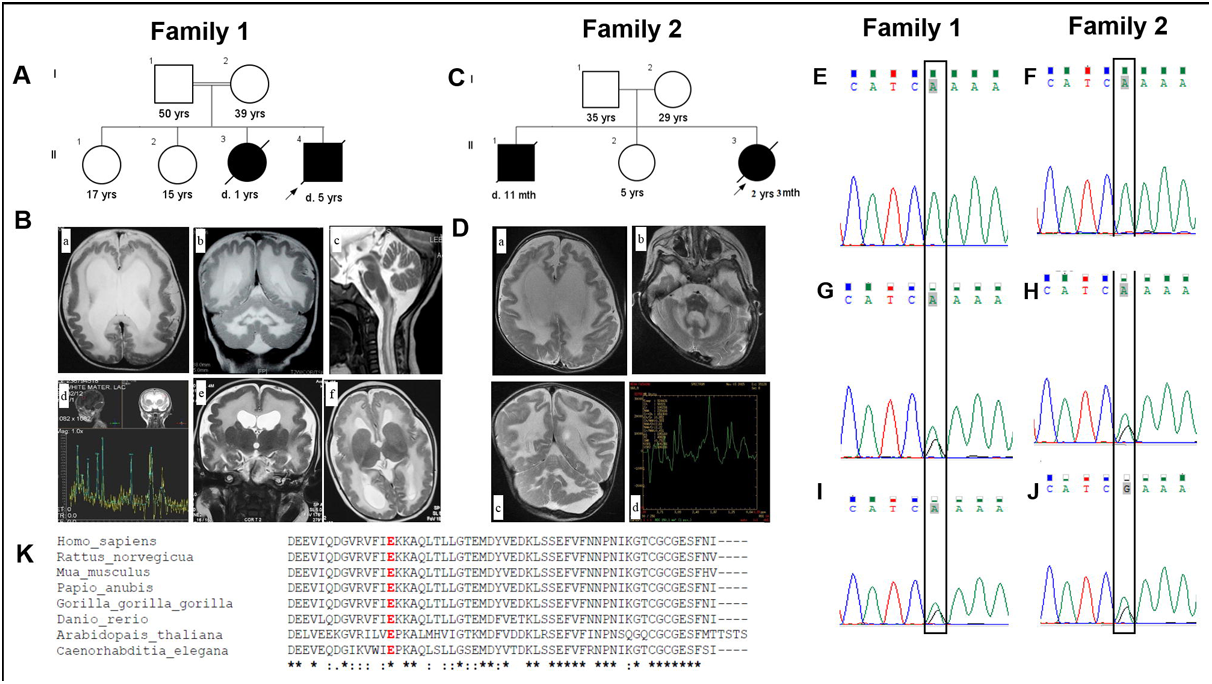
Pedigrees, brain imaging and *ISCA1* variant. (A) Pedigree of Family 1. (B) Brain MRI images of family 1, II-3 and II-4. (a,b) T2 weighted MRI shows pachygyria, moderate dilatation of the ventricles and diffuse T2 hyperintensity in the cerebral and cerebellar white matter and posterior limb of internal capsule in II.3 at 6 months. (c) MRI performed for the same child at 9 months shows myelination abnormality involving the medulla and the cervical cord in the white matter suggestive of delayed myelination and (d) MR spectroscopy shows increased lipid-lactate peaks. (e,f) T2 weighted MRI in II.4 shows pachygyria, moderate dilatation of the ventricles and diffuse T2 hyperintensity at the age of 5 months. (C) Pedigree of Family 2. (D) Brain MRI images of family 2, II-3. (a-c) T2 weighted MRI in II.3 shows diffuse T2 hyperintensity in the cerebral and cerebellar white matter with (d) raised lipid-lactate peaks on MR spectroscopy at the age 1 year 11 months, consistent with dysmyelination. (E-J) Sanger validation of the *ISCA1* variant in family 1 and 2. The pathogenic variation c.259G>A of *ISCA1* is found in homozygous state in (E,F) probands of family 1 and 2 and is heterozygous in their parents (G-J) (K) Comparison of *ISCA1* orthologs from H. sapiens, R. norvegicus, M. musculus, P. anubis, G. gorilla, D. rerio, A. thaliana and C. elegans reveals the high conservation of the Glu87 residue (highlighted in red)

Family 2 was ascertained from our in-house exome data for children with neurodevelopmental disorders. Both II.1 and II.3. (Figure 1C) showed a severe neurodevelopmental disorder with resistant seizures. II.1 had lactic acidosis and white matter disease was documented in medical records of II.1 on computed tomography of brain. II.3 had extensive leukodystrophy involving cerebral and cerebellar white matter with dilated ventricles. Magnetic resonance spectroscopy showed elevated lipid lactate peak in her brain. They succumbed to the disease at the age of 11 months and 2 years 3 months respectively. We note the phenotype observed in all affected subjects in both families is similar to that previously described for all four types of MMDS. Complete clinical details for both families are provided in Table 1. This research work has the approval of the institutional ethics committee. Specific parental consent was obtained for the use of photographs, clinical and research findings for publication.

**Table 1:** Clinical findings observed in our patients.

|  | Family 1 | Family 1 | Family 2 | Family 2 |
| --- | --- | --- | --- | --- |
| Individual | II.3 | II.4 | II.1 | II.3 |
| Origin | Indian | Indian | Indian | Indian |
| Sex | Female | Male | Male | Female |
| Age at assessment | 9 months | 8 months | 7 months | 1 year 11 months |
| Age at demise | 1 year 7 months | 5 years | 11 months | 2 years 3 months |
| Coding DNA change | NA | c.259G>A | NA | c.259G>A |
| Amino acid change | NA | p.E87K | NA | p.E87K |
| Zygosity | NA | Homozygous | NA | Homozygous |
| Exon | NA | 4 | NA | 4 |
| Birth weight (gm/SD) | 3000/-1 | 3600/-1 | 2500/-2 | 2700/-1 |
| **Anthropometry** | - |  |  |  |
| Weight (gm/SD) | 7000/-1 | NA | 6400/-2 | 5800/-6 |
| Length (cm/SD) | 63/-1 | NA | NA | 76/-5 |
| OFC (cm/SD) | 41/-2 | 42/-2 | 42.5/-1 | 43.5/-6 |
| **Neurological abnormalities** |  |  |  |  |
| Neurodevelopmental delay | Severe | Severe | Severe | Severe |
| Feeding difficulty |  | + | + | + |
| Neurologic regression | + | + | + | - |
| Tone | Spasticity | Spasticity | Spasticity | Spasticity |
| Deep tendon reflexes | Exaggerated | Exaggerated | Exaggerated | Exaggerated |
| Strabismus | + | - | - | + |
| Seizures | + | + | + | + |
| Other clinical findings | History of incessant cry, nystagmus | History of incessant cry, tremors in the hands | - | - |
| **MRI findings** |  |  |  |  |
| Pachygyria | + | + | NA | - |
| Cerebral white matter | Leukodystrophy | Leukodystrophy | Leukodystrophy | Leukodystrophy |
| Cerebellar white matter | Leukodystrophy | Leukodystrophy | Leukodystrophy | Leukodystrophy |
| Cerebral ventriculomegaly | + | + | +* | + |
| MRS findings | Elevated lactate peak | Elevated lactate peak | NA | Elevated lactate peak |
| EEG findings | Normal | NA | Abnormal | Normal |
| Ophthalmological findings | Normal VEP and fundus | Stippled pigmentation of fundus | NA | NA |
| Hearing evaluation | NA | NA | Impaired | Normal |
| Blood lactate (mg/dl) | 20.7 | 36 | 65.8 | 40.5 |
| CPK (IU/L) | 132 | 568 | NA | 42 |
| Hematology | Normal | Normal | Normal | Normal |
| Renal function tests | Normal | Normal | Normal | Normal |
| Liver function tests | Normal | Normal | Normal | Normal |
| Metabolic profile** | NA | Normal | NA | NA |

**Abbreviations:** NA: Not available; +: Present; -: Absent; OFC: Occipitofrontal circumference; SD: Standard deviation; MRI: Magnetic resonance imaging; TORCH: Toxoplasmosis, rubella, cytomegalovirus, herpes; VEP: Visual evoked potential; CPK: Creatine phosphokinase; GCMS: Gas Chromatography Mass Spectrometry; TMS: Tandem mass spectrometry; TLC: Thin-Layer Chromatography; FISH: Fluorescence in situ hybridization; RBC: Red blood cells; HPLC: High Performance Liquid Chromatography; \*Observed on computed tomography; \*\*Metabolic profile: serum amino acids, urine organic acids and serum carnitine profiles

Whole exome sequencing (WES) was carried out as described previously to achieve an average coverage depth of 100-130x, such that ~95% of the bases are covered at >20x, with a sensitivity of >90%.^13^ WES raw data was processed using SeqMule and the called variants were annotated with ANNOVAR.^14-16^ Variants were filtered against public databases such as 1000 Genomes Project phase 3, Exome Aggregation Consortium v.0.3.1 (ExAC), National Heart, Lung, and Blood Institute Exome Sequencing Project Exome Variant Server (ESP6500SI-V2) and an in-house database (~139 exomes) and those with a minor allele frequency >1% were excluded. Additionally, variants flagged as low quality or putative false positives (Phred quality score <20, low quality by depth <20) were excluded from the analysis. The overall variant filtering strategy is outlined in Table S1. In view of the consanguinity and phenotype of mitochondrial disorder, we looked for homozygous variants in Family 1. No sequence variants of pathogenic significance were detected in any genes known to be causative for mitochondrial or similar disorders. However, a compelling variant, a non-synonymous missense pathogenic variation, c.259G>A (p.E87K) in *ISCA1* (NM_030940.3) in the homozygous state was found in family 1. The same variant was also observed in proband II.3 of family 2 from the in-house exome sequencing data (Figure S2). The variant was validated by Sanger sequencing in affected subjects from both families (Figure 1E, 1F). Targeted testing of parents using Sanger sequencing confirmed the variation to be heterozygous in them, consistent with an autosomal recessive model of inheritance for this condition (Figure 1G, 1H, 1I, 1J). Unaffected sibs in family 1 were heterozygous carriers of the variant.

The p.Glu87Lys variant of ISCA1 reported here is not present in a homozygous state in 1000 Genomes project, the Exome Variant Server, CentoMD and in our in-house exome data of 139 unrelated individuals from local population. However, it is present in the heterozygous state in 1/118662 individuals (AF = 0.000008427) in ExAC database (60,706 exome dataset). *ISCA1* has a positive Z score (z = 1.29) for missense constraint in ExAC database, which embodies its intolerance to variations. Notably all 17 *ISCA1* rare missense variants listed in ExAC with Allele frequency (AF) < 0.001 are present in heterozygous and none in homozygous state. In the recently made available gnomAD browser comprising of 126,216 exome and 15,136 genome datasets, the variant p.Glu87Lys is observed in heterozygous state in a single individual out of 240,238 (AF = 0.000004163). Finally, this variant was not seen in any other publicly available projects, except ExAC database, which was confirmed via Kaviar v.160204-Public, a database of 162 million SNVs from 35 different projects, underscoring the essential nature of this gene and predicted damaging impact of this variant.

IscA proteins are highly conserved and fundamental to the physiology of prokaryotes and eukaryotes. The human ISCA1 was first identified in a subject with Sjogren’s syndrome as a potential target of autoimmune antibodies.^17^ It was found to be abundantly expressed in human and rat brain and kidneys.^17^ It shares 38% identity and 70% similarity to *E.coli* Isca and similar to its bacterial counterpart.^17;18^ The above reported variant appears to have an impact on the functional Fe-S biogenesis domain of ISCA18 (Figure S1B), at a residue that is highly conserved, as indicated by Clustal Omega multiple sequence alignment (Figure 1K) and ConSurf (Figure 2A, 2B, 2C), with a PhyloP score of 7.442. Additionally we note that the harmful nature of this variation is suggested by its occurrence at residue 87 that lies within a region of 48-90 residues of ISCA1, delineated previously to be instrumental for mediating ISCA1 interaction with IOP1 (iron-only hydrogenase-like protein 1), that plays a role in cytosolic Fe-S protein assembly pathway.^19^ Several functional *in silico* prediction tools were also used to interrogate the damaging consequences of the p.Glu87Lys variation. MutationTaster predicted it to be ‘disease-causing’, similarly the Sorting Intolerant from Tolerant (SIFT) algorithm and Protein Variation Effect Analyzer (Provean) indicate this alteration to be ‘damaging’. The presently reported missense *ISCA1* variation was also predicted to be highly deleterious with a CADD score of 22.5.^20^ This was corroborated by Position Specific Evolutionary Preservation (PANTHER-PSEP) analyses that predicts this SNV to be probably damaging on account of it occurring at an amino acid position estimated to be preserved for ~455 million years. Screening for Nonacceptable Polymorphisms (SNAP2), a web-based tool that combines a variety of information pertaining to evolutionary conservation, secondary structure and solvent accessibility of the protein predicted the p.E87K variant to have a severe detrimental effect with a score of 80 and accuracy of 91%.

**Figure 2:**
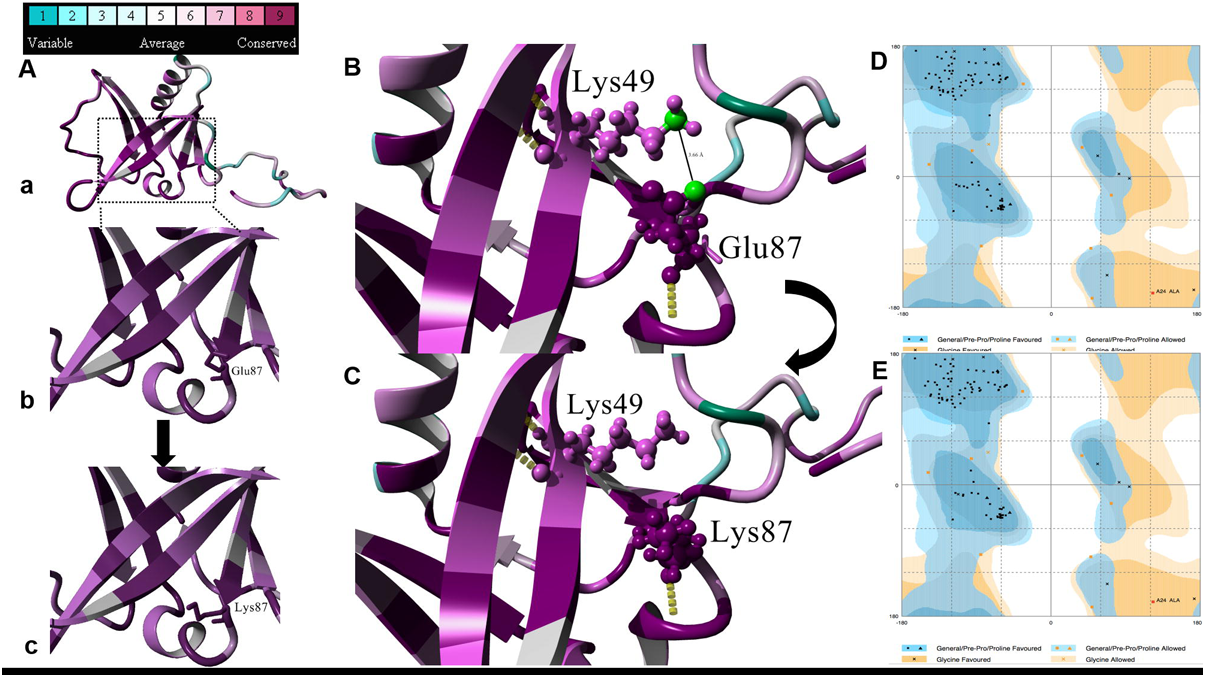
Protein modeling. (A) Protein structure prediction shows replacement of (a,b) the native glutamic acid residue that has an acidic side-chain with (c) lysine bearing a basic side-chain. Color spectrum indicates high conservation of the Glu87 residue. (B) Predicted salt-bridge formation between Glu87 and Lys49 where the distance between the hydrogen donor atom (NZ) of Lys49 and the hydrogen acceptor atom (OE1) of Glu37 is 3.66 Å. (C) Replacement of the Glu87 with mutant Lys87 is predicted to lead to the loss of the salt-bridge between the native residue and Lys49. (D,E) The Ramachandran plots for wild-type and mutant ISCA1. In both (D) wild-type and (E) mutant structures, 90.7% of the all residues were observed in the favoured, 8.4% in the allowed and 0.9% in the outlier regions Note that the favored, allowed and outlier regions are indicated in dark blue, light blue and white.

We used three dimensional (3D) protein structure modeling to assess the structural impact of native acidic Glu87 being replaced by the basic mutant Lys87 residue in the mutant ISCA1. The 3D structure model of wild-type human ISCA1 (UniProt accession: Q9BUE6) and the mutant c.259G>A (p.Glu87Lys) proteins were predicted using SWISS-MODEL web-server. Homology modeling was carried out on the basis of the crystallographic structure of E. coli SufA (sequence similarity of 37.61% to the human ISCA1) that is involved in biosynthesis of Fe-S clusters (PDB ID: 2D2A). The visualization of the protein structure was done using the locally installed YASARA View simulation software.^21^ The Ramachandran plot for native and mutant ISCA1 were generated using RAMPAGE server and were of good quality. In both 90.7% of the all residues were observed in the favored region, with 8.4% in the allowed region and only 0.9% in the outlier region (Figure 2D, 2E). Using Protein Structure Evaluation Suite and Server (PROSESS) that utilizes Volume, Area, Dihedral Angle Reporter (VADAR, the phi (φ) and psi (ψ) backbone dihedral angles for wild-type Glu87 were estimated to be −62.3φ, 138.6ψ and that for mutant Lys87 were −67.8φ, 136.9ψ. The p.E87K alteration has drastic effects on Val50 and Ala90 residues, the hydrogen (H) bonding partners of the native Glu87. In the mutant, the φ angle of Ala90 potentially increases from −101.9φ to −106.2φ, whereas ψ angle decreases from −15.3ψ to −15.1ψ, similarly the φ angle of Val50 decreases from −95.8φ to −95.6φ and ψ angle increases from −125.6ψ to −62.3φ, 138.6ψ. The observed p.E87K variation also seems to affect the main chain residue accessible surface area (ASA) that increases from 41.4 Å2 to 79.9 Å2 and the main-chain residue volume increases from 145.7 Å3 to 178 Å3. Moreover, the side-chain residue ASA increases from 41.4 Å2 to 79.9 Å2 and the solvation energy of the mutant Lys87 decreases from -1.3 kcal/mol to -3.5 kcal/mol. HOPE and ConSurf server based analyses indicated the native Glu87 residue engages in a salt-bridge formation in wild-type ISCA1 with Lys49, where the distance between hydrogen donor (NZ) atom of Lys49 and the hydrogen acceptor (OE1) atom of Glu87 and was found to be 3.66 Å, and this interaction is lost in mutant ISCA1 (Figure 2B, 2C). Additionally, VADAR analysis predicted that the side-chain H bonding interaction of Glu87 with Ser73 and Lys88 are also potentially lost. The neural network based predictor of protein stability, I-Mutant (Reliability Index score 7) and the Eris web-server (score 0.64 ΔΔG kcal/mol) are also consistent in estimating that the p.E87K mutation leads to destabilization of ISCA1 protein. Taken together our structural analysis reflects that the presently reported variation appears to disrupt crucial molecular interactions and reduces the overall ISCA1 protein stability.

Since the two probands are of Indian origin and from the same geographic location (family 1 was consanguineous and family 2 denied any consanguinity) and the same pathogenic variant is identified in both of them, we explored the possibility of a founder effect. After examining the homozygous regions around the *ISCA1* variant in both the probands, generated by the exome sequencing data, we identified only one overlapping region of homozygosity spanning 3.3 Mb in chromosome 9 (Chr9: 85613354-88925774) flanking the variant (Figure S3) suggesting the possibility of a founder mutation in the local population (Table S2 and Table S3). The genes within the shared region are given in Table S4.

To conclude, we describe two independent families, both with two affected children each, with a severe neurodevelopmental disorder associated with a homozygous c.259G>A variant in *ISCA1*. The major clinical features include severe neurodevelopmental delay, seizures, spasticity and regression of milestones. Extensive leukodystrophy, lactate peaks and ventricular dilatation are important changes observed on magnetic resonance imaging of brain. Pigmentary retinopathy, impaired hearing and vision, elevated creatine kinase, lactic acidosis are other likely features of this condition. Taken together, existing literature on Fe-S biogenesis, *in silico* functional analyses, structural modeling predictions, founder effect, the common phenotype in all affected subjects from two families similar to that described in all four types of MMDS that occur due to mutations in genes encoding proteins involved in Fe-S biogenesis, strongly suggest that the presently reported *ISCA1* variant is likely to result another MMDS. Further cases and functional studies would validate our findings.

## Supplemental data

Supplemental data include three figures and four tables.

## Acknowledgement

We thank the families who cooperated with the evaluation of the subjects and consented for participation in this study. This work was supported by US National Institutes of Health funded project titled “Genetic Diagnosis of Heritable Neurodevelopmental Disorders in India: Investigating the Use of Whole Exome Sequencing and Genetic Counseling to Address the High Burden of Neurodevelopmental Disorders” (1R21NS094047-01).

## Web Resources

The URLs for data presented herein are as follows:

Clustal Omega, https://www.ebi.ac.uk/Tools/msa/clustalo/

1000 Genomes Project, http://browser.1000genomes.org/index.html

SWISS-MODEL, https://swissmodel.expasy.org/

RAMPAGE, http://mordred.bioc.cam.ac.uk/~rapper/rampage.php

HOPE, <u>http://www.cmbi.ru.nl/hope/</u>

Exome Variant Server, http://evs.gs.washington.edu/EVS/

The Exome Aggregation Consortium (ExAC) Browser, <u>http://exac.broadinstitute.org/</u>

The Genome Aggregation Database (gnomAD), <u>http://gnomad.broadinstitute.org/</u>

Kaviar, <u>http://db.systemsbiology.net/kaviar</u>

PROSESS, <u>http://www.prosess.ca</u>

VADAR, http://vadar.wishartlab.com/

UniProt, <u>http://www.uniprot.org</u>

ConSurf, <u>http://consurf.tau.ac.il/2016/</u>

OMIM, <u>http://www.omim.org/</u>

MutationTaster, http://www.mutationtaster.org/

The Sorting Intolerant from Tolerant (SIFT), http://sift.jcvi.org/

Protein Variation Effect Analyzer (PROVEAN), http://provean.jcvi.org/

Position-Specific Evolutionary Preservation (PANTHER-PSEP), <u>http://pantherdb.org/tools/csnpScoreForm.jsp</u>

I-Mutant2.0, http://folding.biofold.org/i-mutant/i-mutant2.0.html

Eris, <u>http://dokhlab.unc.edu/tools/eris/</u>

Screening for Non-Acceptable Polymorphisms (SNAP2), https://rostlab.org/services/snap2web/

